# Assembly of sequences of Northern China alligatorweed identifies processes and gene ontologies associated with invasiveness and provides genomic resources for population and gene regulation studies

**DOI:** 10.1101/2023.06.16.545218

**Authors:** David P. Horvath, Yanwen Wang, Fanjin Meng, Mckayla Neubauer, Dasheng Liu

## Abstract

Biological invasions remain a major global challenge. Alligatorweed (Alternanthera philoxeroides (Mart.) Griseb.), native to South America, has had profound negative effects on ecosystem function and economy in Australia, North America, and Asia. It is an invasive and primarily aquatic plant that, despite a documented lack of genetic diversity, is unusually adaptive - thriving in both aquatic and terrestrial environments. However, genetic resources for studying this invasive plant are limited. Here, we have assembled the transcriptome of alligatorweed using all publicly available cDNA sequences. The resulting assembly produced over 500K contigs with an average length of ∼700 bases and an N50 of >1000 bases and contains over 100K probable gene-coding sequences. Although this assembly is slightly smaller than the previously published assembly developed from just cold-treated shoot tips, the new assembly is slightly more complete with over 95% of the conserved plant genes being represented as full length transcripts, and only 2.3% of these conserved genes being unrepresented compared to 2.7% missing in the earlier assembly. Resources from the PANTHER database were used to annotate all transcripts containing long open reading frames. Comparisons to several plant species identified gene ontologies that were over- and under-represented in the alligatorweed transcriptome including cellular transport and cytoskeletal processes and cell signaling, which could explain the high growth rate and phenotypic plasticity that make alligatorweed particularly invasive. We also sequenced and assembled a genomic database for alligatorweed using only short read technologies. This assembly produced over ten million contigs with an average length of only 300 bases and an N50 of 451 bases. However, 88% of the transcripts were represented among the genomic contigs, indicating that these contigs could serve as a source for regulatory elements for genes previously shown to be differentially expressed under various conditions. Kmer analysis indicated that 22% of the alligatorweed genome was comprised of repetitive elements. A similarity search against the plant repetitive element database indicated that long terminal repeat containing elements including copia- and gypsy-like elements made up the bulk of the transposons present in the alligatorweed genome. Additionally, we assembled and annotated a full-length chloroplast and a partial mitochondrial genome. Combined, these resources provide a source of gene sequences that should be useful for more complete genomic assemblies and for investigating gene structure and function in this particularly adaptable and invasive species. The results will provide an excellent starting point for many different investigations into the biology and ecology of alligatorweed, strengthen our understanding of the invasiveness, biology and ecology of invasive plants, and will help develop a reasonable management strategy to reduce risk and costs of the impacts.

## INTRODUCTION

Biological invasions remain a major global challenge (Cohen and Carlton 1998) that has attracted considerable attention from ecologists, policymakers, managers and the public. One species of concern is the Alligatorweed (*Alternanthera philoxeroides*). It is a world-wide invasive weed (Julien and Bourne 1988), and has had profound negative effects on ecosystem function and economy in Australia, China and the United States (Julien and Bourne 1988, Julian et al. 1992, Liu et al. 2017, Buckingham 2002). It primarily infests aquatic ecosystems but can also invade cropland near aquatic regions and reduce crop yields (Julian et al. 1992). Alligatorweed reproduces outside its native range almost entirely through clonal propagation. Stem fragments can easily be broken off and, because they readily float, are distributed along waterways (Sainty et al. 1998). These stem fragments are capable of rooting and establishing new populations.

Alligatorweed is a native of South America, but was spread across several continents by settlers and explorers who utilized it as a fodder crop livestock (Sosa et al. 2008). Alligatorweed has long been present in southern China where it has proven problematic by blanketing water surfaces, choking navigation ways, and damaging local biodiversity and agricultural production (Yin 1992; Ma and Wang 2005; State Environmental Protection Administration of China and Chinese Academy of Sciences2003). Intriguingly, it has recently increased its range into northern Chinese water systems (Liu et al. 2012, Liu et al 2017).

Little is known about the genetic structure of Alligatorweed. It is thought to have several different ploidy levels including tetraploid, pentaploid, and hexaploid varieties (Chen et al. 2015). Due to the clonal nature of its reproduction, there is often little genetic diversity between populations from a given geographical location (Chen et al. 2015, Wang et al. 2005). There have been several RNA sequencing studies (published and unpublished) that are publicly available (Gao et al. 2015, Li et al. 2015, Liu et al. 2019), as well as one published transcriptomic study where the raw data has not been released or made available (Zhu et al. 2015). The genome size of alligator weed ranges from 5.72 to 8.50 pg/2C nucleus or roughly 2.8-4.2 Gb/ haploid genome (Chen et al. 2015)- the largest genome size being indicated for the Chinese variety.

Although full genome sequencing is currently too expensive to obtain a complete genomic sequence for this species, recent work on other non-model invasive weeds has provided a blueprint for obtaining significant amounts of genetic information from small scale and relatively inexpensive short read technologies such as Illumina (Horvath et al. 2018). Such information is invaluable for the identification of genetic differences (markers) needed to follow evolutionary shifts in allele frequency associated with enhanced invasiveness and cold tolerance. Such information is also needed for more general population genetic studies, and for identifying targets and susceptibility to various herbicide or biological control measures employed to reduce alligatorweed impacts on the ecosystem, or to predict potential variants that might be or become resistance to such control measures.

Here, we have obtained all available RNAseq data for the primary variety of alligatorweed endemic to China, and have used this to develop a transcriptome database. This transcript collection was used to inform and annotate a highly fragmented genomic assembly based on less than 20X coverage of the alligatorweed genome. Additionally, this data was used to assemble a complete chloroplast sequence and to develop a comprehensive set of mitochondrial contigs. Our efforts will strengthen our understanding of the invasiveness, ecological adaptation, biology and ecology of this invasive plant, and will also be useful in developing reasonable management strategies to reduce risk and costs of the impacts of this problematic invasive species.

## MATERIALS AND METHODS

### Genomic library construction and analysis

Alligatorweedsplants were collected in Jinan, Shandong, China and then grown under greenhouse conditions. Three individual plants were randomly selected and sent to BGI Global Genomic Services for sequencing. One sample was chosen for library preparation and sequencing. The resulting 150 base paired end reads were trimmed by quality and to remove contaminating primer sequences prior to assembly initially by BGI Global Genomic Services. The resulting reads were subjected to kmer counting (K=17) and assembled using SOAPdenovo by BGI Global Genomic Services.

The assembled database was submitted to NCBI (accession # PRJNA541860) after removal of probable contaminating non-plant sequences. Subsequent analyses were performed on sequences that were further cleaned and trimmed using subroutine in the program BBMAP (Bushnell 2014). BBMAP was used to predict the percentage of highly repetitive sequences within the genome, and the program BlastN was used to identify probable repetitive elements represented in the plant repetitive element database. The program Trinity2.2 (Grabherr et al. 2011) was also used to produce an assembly that, as was found for the perennial invasive weed leafy spurge (Horvath et al. 2018), appeared to be superior to the assembly produced by BGI Global Genomic Services (File S1), however all subsequent analyses were performed on the original assembly produced by BGI Global Genomic Services.

### Plastid assembly

The raw genome sequence was mined for the chloroplast and mitochondrial genomes using the program NOVOPlasty 2.7.2 (Dierckxsens et al. 2017). The large subunit of RUBISCO sequence from alligatorweed (>FR775287.1 Alternanthera philoxeroides chloroplast partial rbcL gene) was used as the initial seed sequence, and the sequence of the Amarantheacea whole chloroplast genome was used as a scaffold (>KX279888.1 *Amaranthus hypochondriacus* cultivar Plainsman chloroplast, complete genome). Likewise, a fragment of an assembled genome sequence that matched a portion of a conserved mitochondrial sequence was used as a seed along with the complete genome sequence of the sugarbeet (*Beta vulgaris*) as a scaffold (>BA000024.1 Beta vulgaris subsp. vulgaris mitochondrial DNA, complete genome). The chloroplast sequence was then annotated initially using the online program DOGMA (Wyman et al 2004) and visualize using the online program GenomeVx (http://wolfe.ucd.ie/GenomeVx/) followed by manual annotation to confirm start and stop codons as well as intron/exon boundaries. Similarly, manual annotation was also performed on the assembled mitochondrial contigs. Both sequences were deposited in genbank (accession #s, MK795965.1 and Y respectively).

### Transcriptome assembly and analysis

The raw transcriptomic sequences were downloaded from all publicly available data from the SRA database at NCBI. These represented four studies-one where stem internode samples were collected from pot-grown plants that had been submerged or not to submerged conditions (PRJNA256237 Yang Ji 2016), one that compared expression differences between plant material (shoot tips, young and mature leaves and mature stems) collected from pot-grown plants in aquatic or terrestrial environments (PRJNA256235) (Gao et al. 2015), one where root samples were collected before and after potassium deprivation (PRJNA263573) (Li et al. 2015), and one where leaf tissue was collected from two different populations following cold-acclimation (PRJNA268359) (Liu et al. 2019). Raw data from the SRA database was downloaded, trimmed as described above. Each separate data set was assembled independently using the program Trinity2.2 (Grabherr et al. 2011). The resulting assemblies were concatenated and clustered to produce a consensus assembly using the program cd-hit-v4.6.6 (Li and Godzik, 2006).

### Assessment of assembly success

The success of the assembly was assessed using the program BUSCO-v3.0 (Waterhouse et al. 2017) in the CyVerse Discovery Environment (Oliver et al., 2013). Surprisingly, an attempt to assemble the combined datasets produced a poorer assembly than assembling each separately-possibly due to differences in library quality and design. However, the clustering of the individual assemblies did produce an assembly superior to the previous best reported assembly (Liu et al. 2019). Long open reading frames (LORFs) were selected using the program Transcript Decoder, in the CyVerse Discovery Environment and annotated by BlastP to the uniport database in CyVerse. The resulting assembly fasta file is available as File S2 and annotations of the assembly in Table S1. BlastP was used to identify transcripts with high similarity to plant transcription factors from the PlantTFDB 4.0 (PlantTFDB-all_TF_pep.fas.gz http://planttfdb.cbi.pku.edu.cn/download.php). Scripts used for cleaning, assembly and annotation are available in File S3.

### Gene ontology studies

The transcripts containing LORFs were blasted against the PANTHER gene ontology database as described by Mi et al. 2013. Redundant orf-containing transcripts from the same gene model (as determined by the Trinity assembly annotation) that mapped to the same PANTHER ontology designation were removed leaving ∼64K annotated genes. The list of genes with associated ontologies were assessed using the PANTHER web-based gene ontology program (Mi et al. 2016).

## RESULTS

### Transcriptome assembly

The transcriptome assembly, following clustering produced 551,955 contigs (Table 1). The completeness of this assembly was quite high as indicated by the representation of conserved plant genes detected using the program BUSCO-v3, with 97.7% of these genes represented in the assembly of which 95.2% were full length. The N50 of the assembly was nearly 1100 bases (Table 1). LORFs were found in 135,146 of these, and of these 106,036 had significant (E-values <0.05) hits to the Uniprot database (Bateman et al. 2018) and 106,149 had significant similarity to the PANTHER protein ontology database (Mi et al. 2019). 55,539 non-redundant gene models mapped to known proteins in the PANTHER gene ontology database and 20,724 different protein ontology identifiers were found. To further assess the representation of the assembly, the peptide sequences from the LORFs were blasted against the arabidopsis TAIR10 peptide database. The results from this analysis indicated that 71% of the transcripts containing LORFs had significant similarity to known arabidopsis genes. Nearly half (15,765) of the arabidopsis genes were represented among the contigs, further indicating the good gene representation among conserved genes in the assembles transcripts. A file containing all clustered transcripts and their annotations can be found in Table S1.

**Table 1.**
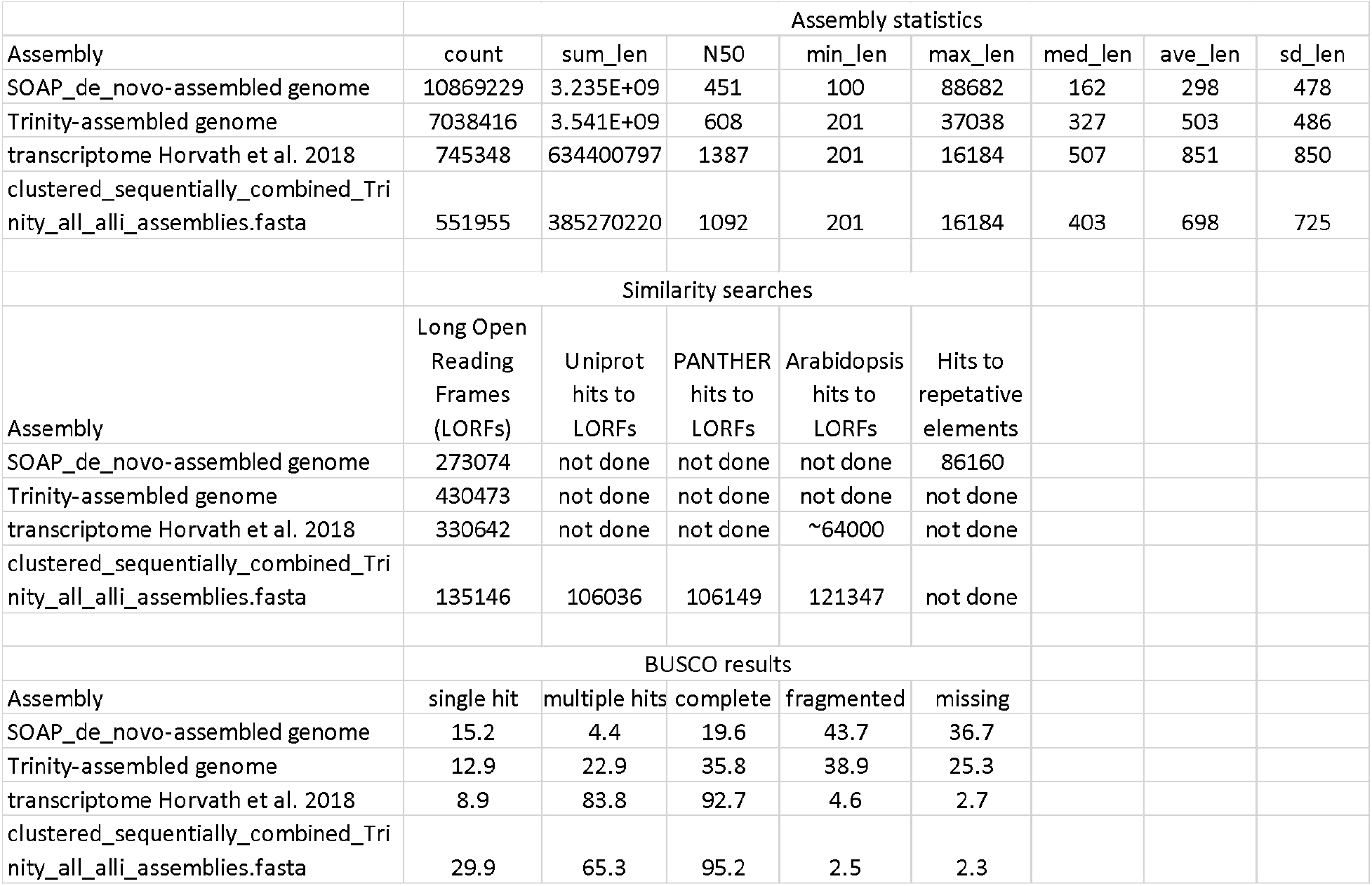
Assembly statistics and analyses.

### Transcriptome contains genes controlling hormone signaling and accumulation and probable transcription factor encoding genes

A list of arabidopsis genes directly involved in hormone metabolism, catabolism, and perception and direct signal transduction was previously generated (Howe et al. 2016). Of the arabidopsis genes with significant similarity to clustered transcripts from the assemblies, 2044 were noted as playing a direct role in hormone accumulation, perception, and signaling (Table S1). Likewise, among the assembled transcripts containing LORFs, 6875 had significant similarity (E<10-20) to previously characterized plant transcription factors in the related species *Amaranthus hypochondriacus* (Table S1). Of these, 920 of the 1259 recognized *A. hypochondriacus* transcription factors were represented with 54 of the 56 transcription factor families present.

### Gene ontology analysis

Given the apparent completeness of the transcriptome, an investigation of over-or under-represented gene classes was warranted. Basic gene ontologies were counted among genes with PANTHER IDs (Figure 1) indicating the presence of all major ontological classes. Ontologies were then compared to five different assembled genomes (arabidopsis (*Arabidopsis thaliana*), sunflower (*Helianthus annuus*), tomato (*Solanum lycopersicum*), cocoa (*Theobroma cacao*), and grape(*Vitis Vernifera*)) (Figure 2,Figure S1 a-e, and Tables S2a-e). There were 136, 74, 72, 81, and 112 significant different ontologies (P,0.00005) arabidopsis, sunflower, tomatoe, cocoa, and grape respectively. Although most differences show a slight under-representation. There are few that are significantly greater in number in most comparisons including axoneme assembly and cilium associated processes -involved in microtubule assembly, cellular signaling, cell transport, protein degradation, and vesicle targeting. Interestingly, the majority are all involved in transport or cell motility and signaling processes. The over-representation of processes controlled by the transcription factor-encoding genes based just on function of similar arabidopsis genes was also examined to look for processes that might be prevalent in the alligatorweed transcriptome. We found 527 different ontologies over-represented (Table S3a). Interestingly, the top process was histone H3-K4 demethylation (GO:0034720). Likewise, a similar assessment examining just hormone-related genes identified 531 significantly over-represented ontologies (Table S3b), with ontologies associated with cytokinin processes (trans-zeatin biosynthetic process (GO:0033466)) being the most significant.

**Figure 1.**
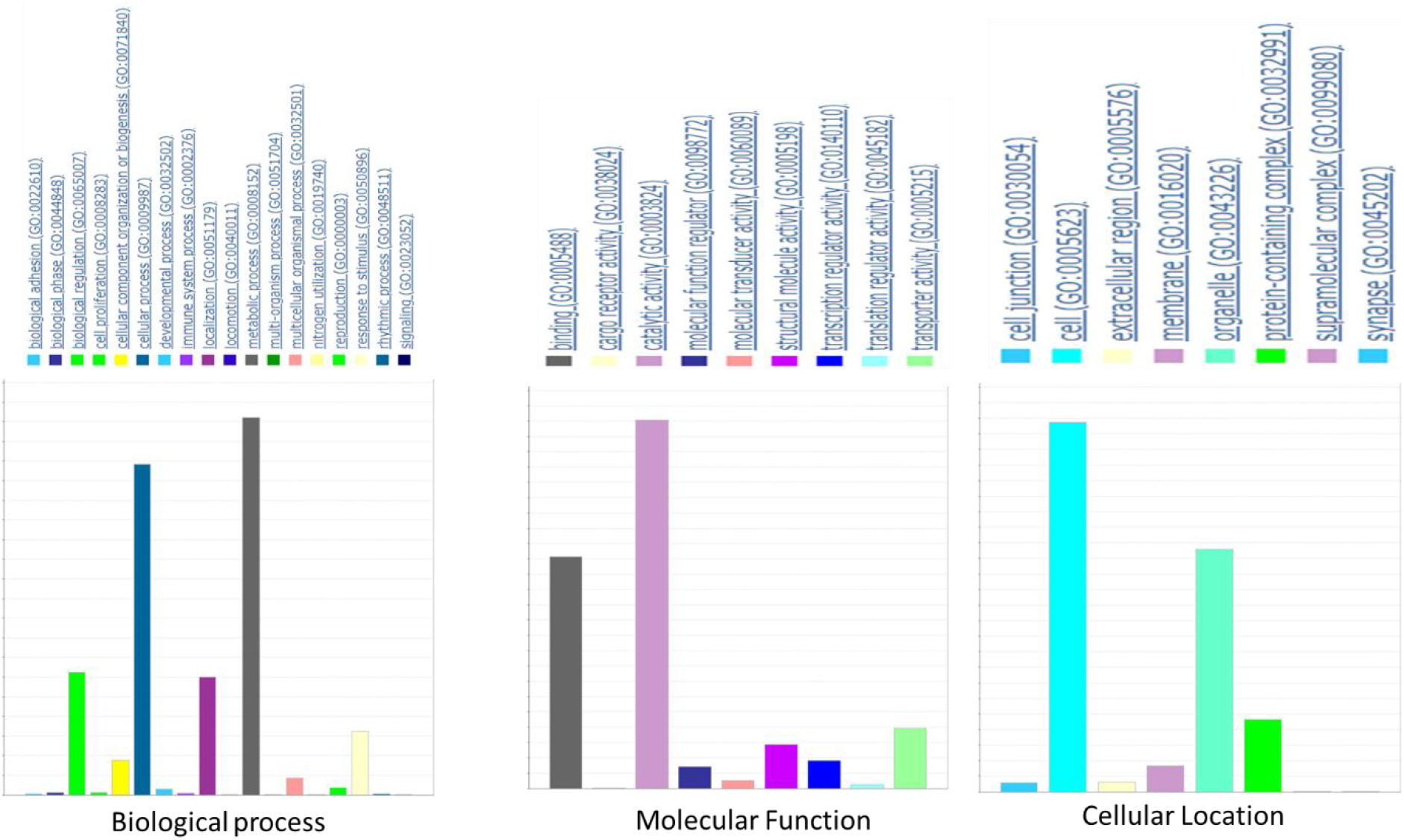
Graphs showing gene ontology counts from all genes with similarity to those in the PANTHER database.

**Figure 2.**
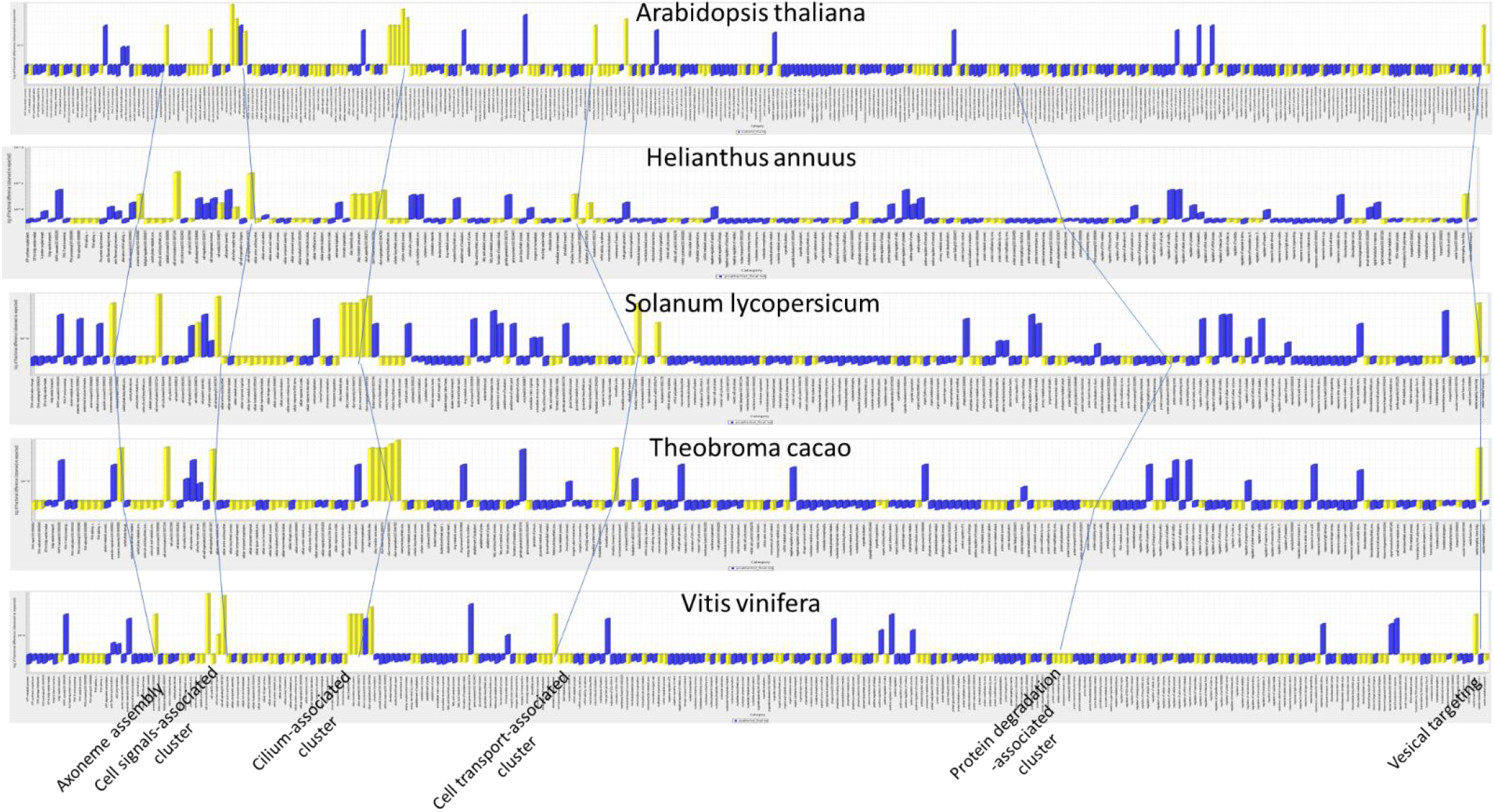
Graphs of over and under-represented gene ontologies when alligatorweed was compared to 5 other species. The yellow bars represent ontologies that were highly significant (p<.000005). The lines connecting bars indicates identical over-represented groups of ontologies with the class noted at the bottom of the chart.

### Genome assembly

From fragments covering only about 16X coverage (based on kmer analysis), we assembled 10,869,229 contigs representing ∼3.2 Gbs-between the minimum and maximum estimated size of the alligatorweed haploid genome (Table 1). Although the longest contig was ∼6.5 times larger than the largest transcript assembled, the N50 value for the contig size of the genomic assembly was less than half the size (N50=451) of the N50 for the transcriptome assembly. We identified 273,074 LORFs within the genomic assembly, of which 83,727 were similar to LORFs found in the transcriptome assembly. This left nearly 190K ORFs that could potentially represent genes not expressed in the plant tissues used to build the sequencing libraries. Thus, this set of ORFs were blasted against the Uniprot database to identify other likely genes missed in the transcriptome assembly. This identified another 10,687 probable functional genes that were missed in the transcriptome assembly, but which are present in the genome assembly, with an additional 401 uniprot annotations that were not present in the transcriptome assembly.

Despite the larger number of LORFS, we found that only ∼63% of the conserved plant genes were represented, and the majority of these were fragmented. However, based on BlastN results, 88% of the assembled transcripts were represented in the assembled genome contigs-suggesting the genome assembly, although fragmented, is reasonably complete regarding the representation of transcribed sequences.

Surprisingly, a kmer analysis using BBMAP indicated a genome size of only ∼1.6 Gbs indicating the probability that the genome likely contains a large number of highly similar sequences which would be expected for an autopolyploid species.

A survey of the assembled genome indicated the possibility of some contamination by sequences associated with whitefly (*Bemisia tabaci*), a common plant pest, based on a large number of sequences from an endosymbiont of whitefly (*Candidatus Hamiltonella defensa* (*Bemisia tabaci*)). Twenty-one genes from this organism were present in the assembly. Several other organisms had relatively high gene representation as well, including another a related species *Candidatus Hamiltonella defensa* (no subspecies designation), *Acetobacter malorum*, *Candidatus Burkholderia schumannianae*, and *Streptococcus agalactiae* CCUG 37742.

### Repetitive elements

Kmer analysis indicated that as much as 22% of the genome might consist of repetitive elements. The program repeat modeler failed to provide output, and thus no de novo repetitive elements were identified. However, we were able to blast the assembly against the known plant repetitive element database to identify probable transposon sequences. Twenty percent of the 61,730 known repetitive elements in the Plant Repetative Element Database were represented in the genomic assembly. Long terminal repeat (LTR) elements including gypsy- and copia-like elements were the most abundant, with RNA and retroelements followed by LINEs and other simple repeats, DNA elements such as En-Spm and Harbringer, with a few non-LTR, satellites, and SINEs with minimal representation (Table 2). Surprisingly only 0.7% of the contigs had hits to repetitive elements. LTR elements were by far the most numerous comprising 69% of the repetitive elements identified.

**Table 2.**
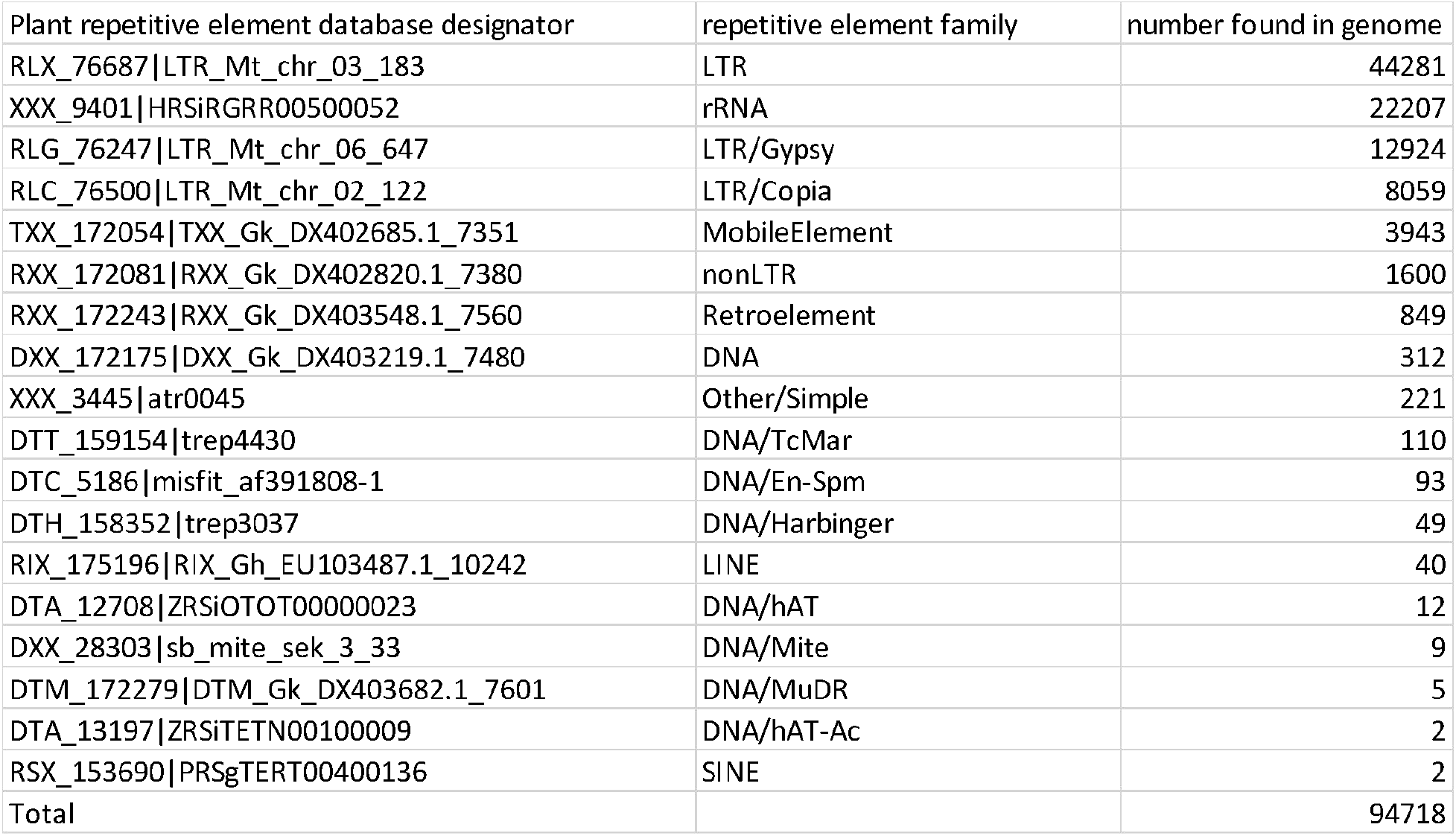
Blast similarity results to Plant Repetitive Element Database.

### Plastid and mitochondrial sequences

A complete chloroplast sequence was assembled from genome sequences and has been submitted to Genbank (accession# MK795965). This resulted in a single circular contig of 152,255 bases (Figure 3). The gene order and evidence of gene editing in specific genes characteristic of the Amaranthaceae family with the most similar sequence being the chloroplast from *Cyathula capitata*. We confirmed the orientation of the direct repeated regions by identifying sequence fragments that contiguously overlapped the ends of the repeat regions. The only significant difference in gene structure was observed in the rpl22 gene. The alligatorweed version of this gene exhibits a slightly truncated open reading frame (lacking the first five amino acids normally found in members of the Amaranthaceae family) due to a mutation in the usual start codon of this gene. The mutation was independently confirmed in cDNA sequences.

**Figure 3.**
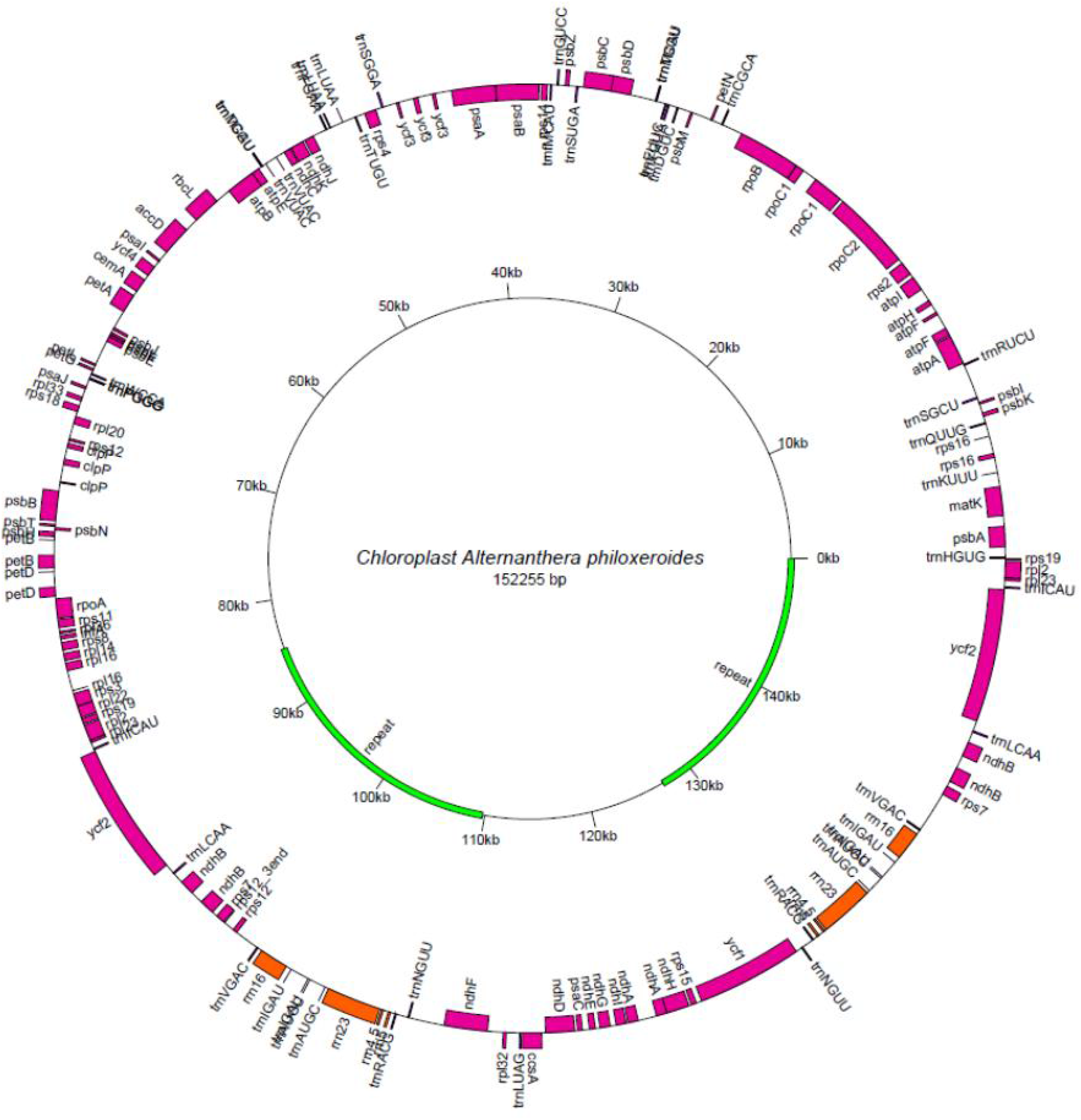
A visual representation of the chloroplast sequence. Protein-coding genes are shown in pink, and ribosomal RNAs are shown in orange. Transfer RNAs are indicated. The orientation of the gene is noted by the position to the outside or inside of the circle. The inner circle indicates the positions of the inverted repeats.

The mitochondrial assembly resulted in three separate contigs covering 289258 bases (supplemental file 1). All well characterized protein coding genes and tRNAs present in related species are included in the discontinuous assembly. The gene order shows regions of syntany with both sugarbeet and quinoa (*Chenopodium quinoa*), but there are notable variations. The majority of the gene editing phenomenon annotated in other related genomes appear to be conserved in the alligatorweed mitochondria. However, the sequence is about 50-100K bases shorter than other related mitochondrial sequences.

## DISCUSSION

### The transcriptome assembly represents a near-complete compliment of alligatorweed genes

We have assembled and annotated the publicly available transcriptome sequences of alligatorweed. A large number of transcripts were assembled, and clustering of these transcripts with a limit of 95% identity resulted in well over 500K different transcripts. The N50 was over 1000 bases, but was less than the N50 (1,387) attained in a previous assembly of just one of the datasets (cold treated leaves samples from northern and southern populations (Liu et al 2019) used for this meta-assembly. The reason why the N50 was shorter from the meta-assembly than from the previously assembled transcriptome is unclear. It is possible that the root samples downloaded from NCBI were of less high quality than the leaves samples and that combining these slightly inhibited the assembly. It should be noted that the length of the longest contig assembled was identical between the previous assembly and this meta-assembly.

Despite the slightly smaller contig sizes, the meta-assembly produced a slightly more complete assembly than the best of the previously assembled transcriptomes of alligatorweed. BUSCO analysis determined that 95.2% of conserved plant genes were represented as complete and full length. This compares to 92.7% in the previous best assembly. Likewise, the meta-assembly resulted in only 2.3% missing conserved plant genes relative to 2.7% missing from the previous assembly. Thus, although marginally shorter in length, this assembly currently represents the best assembly from the alligatorweed transcriptome-particularly when likely unexpressed genes identified in the genomic assembly are included.

It is highly unlikely that there are over 500K genes in alligatorweed, and the fact that only ∼28% of the assembled transcripts contain LORFs suggests that a large amount of the alligatorweed genome is transcribed. A large amount of transcribed non-coding DNA is commonly observed in RNAseq studies like these (Horvath et al 2018 and references therein). The fact that 88% of these transcribed sequences had significant similarity to the assembled alligatorweed genome suggests that the non-coding transcribed sequences are derived from alligatorweed and do not represent poorly assembled contigs or contigs from contaminating species. Thus, these contigs could reasonably serve as a source of additional sequences for developing molecular markers for population genetic studies of this invasive species. However, as was noted in one of the previously published studies of an alligatorweed transcriptome (Horvath et al. 2019), it is likely that a small percentage of these sequences belong to other organisms (viruses, bacteria, fungi, and insects) that are commonly associated with alligatorweed.

Over 88% of the observed LORFs from the transcriptome meta-assembly were similar to previously characterized plant proteins. This suggests that most LORFs represent legitimate alligatorweed genes, and thus are suitable for gene set enrichment analysis and could serve as a source for primer sequences needed to clone and/or investigate the expression patterns of specific genes. The fact that only ∼22% of these are non-redundant is remarkable and suggests that there may be a large number of transcripts from any given gene. The alligatorweed genome is quite large, and the ploidy level could be as high as 6X (Chen et al. 2015). Thus, the high number of redundant transcripts could be due to multiple homeologous genes in the clustered consensus sequence. Although there appears to be evidence of relatively little heterozygosity in the alligatorweed genome (Chen et al. 2015), redundant transcripts could also represent allelic variation as well as alternately spliced transcripts. Also, although BUSCO-v3 analysis and the high N50 values of the transcriptome assembly suggests that the transcriptome primarily represents full length coding sequences, some transcripts are undoubtedly fragmented. A fragmented assembly would also drive the number of assembled contigs higher than the number that is actually present in the alligatorweed genome. These caveats aside, it is likely that we have assembled at least 20K non-redundant alligatorweed genes with an average length of over 1,400 bases. This dataset should provide an excellent starting point for many different investigations into the biology and ecology of alligatorweed.

### The genome assembly provides a rich source of structural genes and their potential regulatory regions

The genomic assembly produced over 10 million small contigs and scaffolds. However, the total length of the assembly was over 3.2 Gb in size- and thus may well represent a majority of the alligatorweed genome which has been estimated to be 2.8-4.2 Gb/ haploid genome (Chen et al. 2015). The fragmented nature of the assembly is typical of assemblies produced from only short-read technologies such as Illumina (Yao et al. 2012). It is possible that scaffolding the assembly with a reference sequence from a related species such as grain amaranth (*Amaranthus hypochondriacus* L.) or sugarbeet could improve the assembly. Likewise, this assembly would benefit from additional sequencing with a long-read technology such as PacBio (Rhoads and Au 2015) or Nanopore methods (Branton and Dreamer 2019) or even the production and sequencing of jump-libraries using short-read technologies.

In this assembly, 88% of transcripts were represented among the genomic contigs. Additionally, there were more than 10K additional assembled LORFs that had similarity to known proteins that were not represented in the transcriptome assembly. Since many genes are only induced during specific times in development or following exposure to specific environmental conditions, it is unlikely that any transcriptome assembly would contain a full complement of genes. Thus, these LORFs could serve as a rich source for non-housekeeping genes including those required for unique developmental processes or responses to specific environmental or biotic stresses.

The fact that there are numerous coding sequences present in the genome assembly suggest that it could also be used to identify promoter sequences from genes that might be differentially regulated as those noted by Horvath et al. (2018) between cold treated alligatorweed populations collected from different regions of China, or other future studies. Similar methods successfully identified promoters in the fragmented genome assembly of another invasive weed leafy spurge (Horvath et al. 2018). These promoter sequences could be used to look for over-represented sequences characteristic of transcription factor binding sites among coordinately regulated genes.

### Assembly of plastid genomes confirms similarity to other related plant species

Using a reference scaffolding method, we were able to assemble a likely full-length chloroplast genome sequence as well as a slightly fragmented but likely near full-length mitochondrial sequence from the alligatorweed genomic sequence. The whole chloroplast sequences was slightly larger (152255 bp) than that of chloroplast of amaranth (151461 bp). The gene order and most of the characteristic start and stop codons, including unusual start codon usage commonly found in other members of the Amaranthaceae were also found in the assembled plastid sequences of alligatorweed. Only one structural gene (rpl22) was significantly altered due to a mutation that impacted the normal start codon of this gene. Likewise, the assembled mitochondrial sequence was 289258 bp, slightly shorter than the 315003 bp long sequence of quinoa mitochondrial genome and much smaller than the mitochondrial genome of the sugarbeet sequence (501020 bp) used to guide the assembly. The initial assembly contained significant chloroplast sequences which were removed, and also originally contained a long region that was repeated in two of the contigs but which was deleted from one. It is possible that some of these sequences may actually be part of the alligatorweed mitochondrial genome. However, without evolutionary support in the form of similar repeats or chloroplast-like sequences in related species, we chose to remove them from our assembly. Long read technologies will be needed to resolve these issues. However, again, regions of syntenic gene order was observed within the three assembled contigs, but there some clear differences observed. For example, the ATP6 gene appears to be located between the tRNA-Pro and the rps7 gene in alligatorweed rather than between tRNA-His and tRNA-Met genes as was observed for both quinoa and sugarbeet. The fact that no single contig was assembled for the mitochondria is not terribly surprising. Plant mitochondria contain several repeated regions, and recombination events between these repeats are well documented. These recombination events can lead to multiple genome conformations within the same organism and even result in multiple subunits forming smaller circular fragments (Burger et al. 2003).

Plastid and mitochondrial sequences are often used for population genetic analyses. These sequences should provide an excellent source for developing primers specifically designed to variable regions and thus facilitate evolutionary relationships between alligatorweed population within and outside of China. Additionally, our previous study highlighted differences in the expression of photosynthetic genes between the population that recently extended its range northwards and that of the more central population (Liu et al. 2019). Thus, these plastid and mitochondrial sequences could be used to look for allelic changes associated with the northward range extension in this species. Also, alligatorweed has been shown to have an unusually high photosynthetic capability (Liu et al. 2007). Thus, these sequences could also provide some insight into photosynthesis and requisite metabolic activity in alligatorweed.

### The genome assembly identifies other species associated with alligatorweed

The possible contamination from other species is clearly not limited to the transcriptome assembly. Indeed, a significant number of genes belonging to an endobacteria of white fly (*Bemisia tabaci*) (Rao et al. 2012), as well as those of related species of endobacteria were assembled from the whole genome sequence data. This information provides a strong indication that the alligatorweed used to create the genomic libraries were infested with this common plant pest. However, this information is of interest since it identifies other species that are in close association with the alligatorweed tissues collected, and could provide information on possible biocontrol agents or other important intraspecies ecological associations involving alligatorweed.

### Repetitive elements are indicative of a flexible genome

Kmer analysis indicated as much as 22% of the alligatorweed genome is comprised of repetitive sequences. Although this could not be confirmed by analysis of the genome assembly using the program repeatmodeler. Blast analysis with the known plant repetitive element database identified several characteristic transposable element families. However, very few of the contigs appeared to contain previously characterized repetitive elements. This suggests that a large number of novel elements may exist in the alligatorweed genome, and that further study is warranted. Alligatorweed is known to have highly plastic growth, and is capable of growing equally well in both aquatic and terrestrial environments. This is despite the fact that there is relatively little genetic diversity among the invasive biotypes. Thus, the presence of transposable elements could provide some of the necessary diversity that allows alligatorweed to be particularly adaptive and invasive. It will be interesting to look for evidence of transposable element movement in different populations of alligatorweed – particularly in documented cases of changes to invasive behavior as noted in the recent range expansion into more northern regions of China or in response to terrestrial vs aquatic environments.

### Gene ontology analysis

Genes involved in cellular transport and cytoskeletal processes and cell signaling are over-represented in the alligatorweed genome relative to the number of genes with similar ontologies in 5 other plant species. Alligatorweed is known to be a very fast-growing plant that can double its biomass every 41 days (Julian et al. 1992) and increase its area by 200% annually (Clements et al. 2011). The rapid growth potential of alligatorweed could be the reason for over-representation of ontologies associated with transport, since metabolites would need to be rapidly moved from source tissues to sink tissues as the plant grows. The role of the cytoskeleton is also well recognized in cell division processes, and thus might explain the over-representation of genes with this ontology. It was interesting to note that the number of significant differences in gene ontologies increased as the phylogenetic distance between the compared species. This would be expected, and indicates that the differences observed are likely real and potentially meaningful.

The observation that cytokinin-related ontologies were among the most significantly over-represented when just the hormone-related genes were examined is also of interest. Cytokinin, together with auxin are the primary hormones associated with growth and development in plants (Moubayidin et al 2009). Cytokinin is associated with positively regulating CYCLIN D type genes required for cell division (Riou-Khamlichi et al. 1999). The observation that genes involved with cytokinin production and signaling are over-represented in the alligatorweed genome would be consistent with the observed high growth rate of this species.

The observation that among the genes encoding probable transcription factors, that histone H3-K4 demethylation was the most over-represented process is also of interest. Alligatorweed is known to be fairly lacking in genetic diversity (Chen et al. 2015). However, its considerable plasticity in its response to different environments, and ability to grow in both aquatic and terrestrial ecosystems were noted above. It has been proposed that the physiological plasticity of alligatorweed is likely due to its ability to regulate genes epigenetically (Geng et al. 2013). The observation of histone demethylation as a significantly over-represented process would be consistent with a high level of, or at least a high capability for epigenetic modification.

### Conclusion

The meta assembly of the alligatorweed transcriptome and the fragmented assembly of the alligatorweed genome represents the current best assemblies available for this agronomically and environmentally important invasive plant species. These resources will provide a rich resource for sequences of structural genes and provide sequences that could be leveraged to explore gene regulation and evolution of this surprisingly adaptable species.

Additionally, gene set enrichment analysis of the transcriptome, and of specific genes with similarity to known hormone action and gene regulation provide observations regarding unique aspects of alligatorweed growth and development that may provide clues to its unusual prolific and plastic growth. Genes involved in cellular transport and cytoskeletal processes and cell signaling are over-represented, the rapid growth potential of alligatorweed could be the reason for over-representation of ontologies associated with transport.The identification of specific transposable elements and evidence of increased chromatin modification should also allow investigations into any associated genome flexibility that could also be related to its recent northward range expansion and/or its phenological plasticity. This dataset should provide an excellent starting point for many different investigations into the invasiveness, biology and ecology of alligatorweed.Finally, this work also points towards the need to further develop genomic sequences-particularly using long-read technologies to develop a needed comprehensive genome sequence. A full genome sequence would provide even greater abilities to explore the evolution of this remarkable albeit problematic species.

chart into the template, be sure it is already sized appropriately and paste it immediately after the and figures. Tables should be simple and concise. It is preferable to use the Table Tool in your word-processing package, placing one entry per cell, to generate tables.

## Supporting Information

File S1: The genomic fasta file produced by Trinity2.2.

File S2: Clustered transcriptome fasta file produced by Trinity2.2.

File S3: All scripts used to clean, trim and assemble the genomic and transcriptomic data.

Table S1: Annotated transcript list from various similarity searches of the transcriptome assembly.

Table S2a-e: Tables showing the results of a gene ontology enrichment analysis in a pairwise comparison of transcription factors-encoding genes for arabidopsis (a) sunflower (b) tomato (c) cocoa (d) and grape (e) as compared between alligatorweed as analyzed with using a binomial pair-wise comparison with the online tool from PANTHER.

Table S3a-b: Tables showing the results of a gene ontology enrichment analysis in a pairwise comparison of transcription factors-encoding genes (a) and of hormone-associated genes (b) as compared between alligatorweed and arabidopsis as analyzed with using a binomial pair-wise comparison with the online tool from PANTHER.

Figures S1a-e: Large PNG files showing over- and under-represented gene ontology classes from alligatorweed relative five other species. Representation was determined by binomial distribution of ontological classes pairwise between alligatorweed and each of the other species individually.

## AUTHOR INFORMATION

### Author Contributions

The manuscript was written through contributions of all authors. All authors have given approval to the final version of the manuscript. Plant material and sequencing was collected by Prof. Liu and associates, bioinformatic analyses were completed by Dr. Horvath and associates.

### Funding Sources

This work was supported by 2016 Shandong Xiaoqing River & Nasi Lake Biodiversity Program and 2018 Shandong Xiaoqing River Basin Aquatic Biodiversity Survey Program.

## ABBREVIATIONS

NCBI: National Center for Biotechnology Information
PANTHER: Protein Analysis Through Evolutionary Relationships.

## Notes

### Competing Interest Statement

The authors have declared no competing interest.

## REFERENCES

Bateman A, Martin M-J, Orchard S, Magrane M. Alpi E, Bely B, et al. 2018. UniProt: a worldwide hub of protein knowledge. Nucleic Acids Res 47:D506–D515.

Branton and Dreamer 2019 The Development of Nanopore Sequencing. In Eds Branton and Dreamer, Nanopore Sequencing. Pp 1–16. WORLD SCIENTIFIC doi:10.1142/9789813270619_0001.

Buckingham GR. 2002. Alligator weed. In Biological Control of Invasive Plants in the Eastern United States; Driesche, R. V., Blossey, B., Hoodle, M., Lyon, S., Reardon, R., Eds.; USDA Forest Service: Washington, DC, 2002; pp 5−15.

Burger G, Gray MW, Lang BF. 2003. Mitochondrial genomes: anything goes. Trends in Genetics 19: 709–716.

Bushnell B. 2014. BBMap: a fast, accurate, splice-aware aligner. Report Number: LBNL-7065E, Lawrence Berkeley National Laboratory, Berkeley, CA.

Chen Z-Y, Xiong Z-J, Pan X-Y, Shen S-Q, Geng Y-P, Xu C-Y, Chen J-K, Zhang W-J. 2015. Variation of genome size and the ribosomal DNA ITS region of Alternanthera philoxeroides (Amaranthaceae) in Argentina, the USA, and China. Journal of Systematics and Evolution 53: 82–87.

Cohen AN, Carlton JT. 1998. Accelerating invasion rate in a highly invaded estuary. Science 279: 555−558.

Clements D, Dugdale TM, Hunt TD. 2011. Growth of aquatic alligator weed (Alternanthera philoxeroides) over 5 years in south-east Australia. Aquatic Invasions 6: 77–82. doi: 10.3391/ai.2011.6.1.09.

Dierckxsens N, Mardulyn P, Smits G. 2017. NOVOPlasty: de novo assembly of organelle genomes from whole genome data. Nucleic Acids Res. 45: p. e18

Gao L, Geng Y, Yang H, Hu Y, Yang J. 2015. Gene expression reaction norms unravel the molecular and cellular processes underpinning the plastic phenotypes of Alternanthera philoxeroides in contrasting hydrological conditions. Front. Plant Sci. 6: 991. 10.3389/fpls.2015.00991

Geng Y, Gao L, Yang J. 2013. Epigenetic flexibility underlying phenotypic plasticity. Prog Bot. 74:153–163.

Grabherr MG, Haas BJ, Yassour M, Levin JZ, Thompson DA, Amit I, Adiconis X, Fan L, Raychowdhury R, Zeng Q. 2011. Full-length transcriptome assembly from RNA-Seq data without a reference genome. Nature Biotechnology 29: 644–652.

Horvath DP, Patel S, Doğramaci M, Chao WS, Anderson JV, Foley ME, Scheffler B, Lazo G, Dorn K, Yan C, Childers A, Schatz M, Marcus S. 2018. Gene Space and Transcriptome Assemblies of Leafy Spurge (Euphorbia esula) Identify Promoter Sequences, Repetitive Elements High-Quality Markers, and a Full-Length Chloroplast Genome. Weed Science 66: 355–367.

Howe GT, Horvath DP, Dharmawardhana P, Priest HD, Mockler TC, Strauss SH. 2015. Extensive transcriptome changes during natural onset and release of vegetative bud dormancy in Populus. Frontiers in Plant Science 6: 00989. DOI=10.3389/fpls.2015.00989.

Li L, Xu L, Wang X, Pan G, Lu L. 2015. De novo characterization of the alligator weed (Alternanthera philoxeroides) transcriptome illuminates gene expression under potassium deprivation. J. Genet. 94: 95–104.

Li W, Godzik A. 2006. Cd-hit: a fast program for clustering and comparing large sets of protein or nucleotide sequences. Bioinformatics 22: 1658–1659.

Liu D, Zhang X, Zhou J, Zhang B. 2007. A preliminary study on diurnal courses of photosynthesis in an invasive species, Alternanthera philoxeroides, North China. Chinese Environmental Pollution & Control; 2007-09

Liu DS, Hu JF, Horvath DP, Zhang XJ, BianXY, Chang GL, Sun XH, Tian J. 2012. Invasions and impacts of Alternanthera philoxeroides in the upper Xiaoqing River Basin of Northern China. Journal of Aquatic Plant Management 50: 19–24.

Liu D, Wang R, Gordon DR, Sun X, Chen L, Wang Y. 2017. Predicting Plant Invasions Following China’s Water Diversion Project. Environ. Sci. Technol. 51: 1450−1457.

Liu D, Horvath D, Li P, Liu W. 2019. RNA sequencing characterizes transcriptomes differences in cold response between northern and southern Alternanthera philoxeroides and highlight adaptations associated with northward expansion. Frontiers in Plant Science.10:24.

Julien MH, Bourne AS. 1988. Alligator weed is spreading in Australia. Plant Protection Quarterly 3: 91–96.

Julien M, Bourne A, Low V. 1992. Growth of the weed Alternanthera philoxeroides (Martius) Grisebach, (alligator weed) in aquatic and terrestrial habitats in Australia. Plant Protection Quarterly 7: 102–108.

Ma RY, Wang R. 2005. Invasive mechanism and biological control of alligator weed Alternanthera philoxeroides(Amaranthacae) in China. Chinese J.Appl. Environ. Biol. 11: 246–250.

Mi H, Huang X, Muruganujan A, Tang H, Mills C, Kang D, Thomas PD. 2016. PANTHER version 11: expanded annotation data from Gene Ontology and Reactome pathways, and data analysis tool enhancements. Nucl. Acids Resdoi: 10.1093/nar/gkw1138.

Mi H, Muruganujan A, Casagrande JT, Thomas PD. 2013. Large-scale gene function analysis with the PANTHER classification system. Nature Protocols 8: 1551–66. doi:http://dx.doi.org/10.1038/nprot.2013.092.

Mi H, Muruganujan A, Huang JX, Ebert D, Mills C, Guo X, Thomas PD. 2019. Protocol Update for large-scale genome and gene function analysis with the PANTHER classification system (v.140) Nat Protoc. 14: 703-721. doi: 10.1038/s41596-019-0128-8.

Moubayidin L, Di Mambro R, Sabatini S. 2009. Cytokinin-auxin crosstalk. Trends Plant Sci. 14: 557–62.

Oliver SL, Lenards AJ, Barthelson RA, Merchant N, McKay SJ. 2013. Using the iPlant collaborative discovery environment. Current Protocols in Bioinformatics 1: 22.

Rao Q, Wang S, Su Y-L, Bing X-L, Liu S-S, Wang X-W. 2012. Draft Genome Sequence of “Candidatus Hamiltonella defensa,” an Endosymbiont of the Whitefly Bemisia tabaci. Journal of Bacteriology 194: 3558. DOI: 10.1128/JB.00069-12.

Riou-Khamlichi C, Huntley R, Jacqmard A, Murray JAH. 1999. Cytokinin activation of Arabidopsis cell division through a D-type cyclin. Science 283: 1541–1544.

Rhoads A, Au KF. 2015. PacBio sequencing and its applications. Genomics Proteomics Bioinformatics 13: 278–289.

Sainty G, McCorkelle G, Julien M. 1998. Control and spread of alligator weed Alternanthera philoxeroides (Mart.) Giseb., in Australia: lessons for other regions. Wetlands Ecol. Manage., 5: 195–201.

Sosa AJ, Telesnicki M, Cardo V, Telesnicki MC, Julien MH. 2008. The evolutionary history of an invasive species: Alligator weed, Alternanthera philoxeroides. In: Julien MH, Sforza R, Bon MC, Evans HC, Hatcher PE eds. Proceedings of the XII International Symposium on Biological Control of Weeds. Wallingford: CABI. 435–442.

State Environmental Protection Administration of China and Chinese Academy of Sciences, 2003, The Announcement About the First List of Invasive Species in China.

Wang BR, Li WG, Wang J. 2005. Genetic diversity of Alternanthera philoxeroides in China. Aquatic Botany 81: 277–283.

Waterhouse RM, Seppey M, Simao FA, Manni M, Ioannidis P, Klioutchnikov G, Kriventseva EV, Zdobnov EM. 2017. BUSCO applications from quality assessments to gene prediction and phylogenomics. Mol Biol Evol. 35: 543–548.

Yao G, Gao H, Minx P, Warren WC, Weinstock GM. 2012. Graph accordance of next-generation sequence assemblies. Bioinformatics 28: 13–16.

Yin RG. 1992. The occurrence and hazard of Alternanthera philoxeroides in vegetable land. Chinese Weed Sci. 1:13.

Zhu Z, Zhou C, Yang J. 2015. Molecular phenotypes associated with anomalous stamen development in Alternanthera philoxeroides. Front Plant Sci 6:242.

